# Construction of high-density genetic maps and detection of QTLs associated with Huanglongbing infection in citrus

**DOI:** 10.1101/330753

**Authors:** Ming Huang, Mikeal L. Roose, Qibin Yu, Dongliang Du, Yi Zhang, zhanao Deng, Michael Irey, Ed Stover, Fredrick G. Gmitter

**Author notes:** Corresponding author Fredrick G. Gmitter Jr.

## Abstract

No true resistance to Huanglongbing (HLB), a citrus disease associated with infection of *Candidatus* Liberibacter asiaticus (*C*Las), is found within commercial citrus cultivars, though trifoliate orange (*Poncirus trifoliata*) has been described as resistant or tolerant. Through genotyping an intergeneric F_1_ population by Genotyping-by-Sequencing, high-density SNP-based genetic maps were constructed separately for trifoliate orange and sweet orange (*Citrus sinensis*). Both genetic maps exhibited high synteny and high coverage of citrus genome. After exposure to intense HLB pressure for two years, Ct value of qPCR for *C*Las detection in leaves throughout ten time points during the next three years was above 35 in trifoliate oranges, under 28 in sweet oranges, and ranged from 24 to 38 and exhibited obvious segregation among progenies. Phenotypic data of percentage of healthy trees showed high correlation with the Ct value. By mapping the two traits at all time points, a total of nine clusters of QTLs were detected, of which five, respectively located on LG-t7 and LG-t8 of trifoliate orange map and LG-s3, LG-s5 and LG-s9 of sweet orange map, collectively explained a major part of the phenotypic variation. This study provides a starting point for citrus breeding to support long-term control of this devastating disease.

**Highlight:** 1). Constructed the first high-density genetic map for trifoliate orange (*Poncirus trifoliata*)

2). The first report on identification of QTLs related to Huanglongbing in citrus.

**Abbreviations:**

ACP: Asian citrus psyllid
*C*Las: *Candidatus* Liberibacter asiaticus
cM: centiMorgans
Ct: Cycle threshold
HLB: Huanglongbing
IM: Interval mapping
KW: Kruskal-Wallis
LG: Linkage group
LOD: Logarithm of odds
QTL: Quantitative trait locus
RAD: Restriction site associated DNA
rMQM: restricted multiple QTL mapping
SNP: Single nucleotide polymorphism.

## Introduction

Huanglongbing (HLB), commonly known as citrus greening, is the most devastating disease in citrus plantations worldwide. Since the first identification of the disease in Florida in 2005, HLB has spread throughout the state, and is now found in most states where citrus is grown. Due to widespread infection and lack of effective management strategy, Florida’s nearly 11 billion dollar citrus industry has experienced a rapid and continuous decline. The production of sweet orange dropped from 240 million boxes in 2004 to 45 million in 2018 (Florida Citrus Commission: http://www.floridacitrus.org).

HLB is generally considered to be caused by three species of *Candidatus* Liberibacter, of which *Candidatus* Liberibacter asiaticus (*C*Las) is the most widespread and virulent species and is the only species reported in citrus industry of United States. *C*Las is a heat-tolerant gram-negative bacteria, resides only in the phloem of plant hosts, and is vectored by the sap-sucking Asian citrus psyllid (ACP, *Diaphorina citri*) (Wang *et al.*, 2017a). As an obligate and insect-transmitted plant pathogen, *C*Las attacks all species and hybrids in the genus of *Citrus* as well as some closely related genera (Ramadugu *et al.*, 2016). Species infected include sweet orange (*Citrus sinensis*), mandarin (*Citrus reticulata*), pummelo (*Citrus maxima*), citron (*Citrus medica*), grapefruit (*Citrus paradisi*), lemon (*Citrus limon*), lime (*Citrus aurantifolia*), kumquat (*Citrus japonica*), and orange jasmine (*Murraya paniculata*). Most commercial citrus cultivars are known as highly susceptible to HLB. Within cultivated citrus resistance or tolerance to HLB was only found in some varieties that are commonly used as rootstocks, such as trifoliate orange and some of its hybrids (Albrecht and Bowman, 2012).

Until now very little is known about the molecular mechanism of pathogenesis following the infection of citrus by *C*Las (Martinelli and Dandekar, 2017). As to the genetic architecture of citrus resistance or susceptibility to HLB, there are no compelling progress, and no QTLs related to *C*Las infection and HLB tolerance response are reported in citrus. Moreover, so far there is no sustainable management and control of HLB where it is endemic. Breeding and development of HLB resistant or tolerant cultivars is widely regarded to be the most practical strategy to support long-term control of this severe disease in the field. The reports of variability for resistance or tolerance to HLB within citrus and its relatives encourages breeding and selection for resistant or tolerant genotypes (Killiny *et al.*, 2017; Miles *et al.*, 2017; Ramadugu *et al.*, 2016; Richardson and Hall, 2013). Traditional breeding through crossing elite cultivars with resistant materials can achieve this objective. However, the introgression of resistant germplasm into elite cultivars will likely require multiple rounds of backcrossing to recover desirable commercial traits. In addition, the breeding cycle for citrus ranges from 5 to 10 years, and the rescue of the citrus industries through HLB-resistant or tolerant cultivars demands considerable urgency. Identification of QTLs associated with resistance or tolerance to HLB in citrus can facilitate more rapid development of resistant cultivars through marker-assisted selection or genome engineering.

To identify QTLs associated with phenotypic traits, a genetic map with high resolution and fine accuracy is crucial. Nowadays, the wide application of high-throughput genome sequencing and efficient SNP genotyping allows the construction of genetic maps with numerous markers at an acceptable cost (Deschamps *et al.*, 2012). While initially confined to herbaceous plants, high-density genetic mapping is increasingly extended to perennial woody plants. Using high-throughput genotyping, the saturation of genetic maps has been greatly improved for some citrus species, such as sweet orange, mandarin and pummelo (Curtolo *et al.*, 2017; Guo *et al.*, 2015; Imai *et al.*, 2017; Lyon, 2008; Ollitrault *et al.*, 2012a; Shimada *et al.*, 2014; Yu *et al.*, 2016). However, in comparison with well-studied model and agronomic plants, citrus lags in development of high-density, high-resolution genetic maps with fine accuracy and precision. Moreover, so far there is no high-density genetic map for trifoliate orange.

This study inaugurates investigation of *C*Las infection and HLB tolerance in citrus by repeated phenotyping of segregating populations and QTL mapping. The objectives of this study were: 1) to evaluate status of *C*Las infection through two phenotypic traits, Ct value of *C*Las detection and percentage of healthy trees, among a field population exposed to intense HLB pressure; 2) to construct high-density genetic maps separately for trifoliate orange and sweet orange through genotyping a mapping population using Genotyping-by-Sequencing; 3) to identify QTLs associated with *C*Las infection using the phenotypic data separately in trifoliate orange and sweet orange genetic maps.

## Materials and Methods

### Plant materials

Genotyping was carried out in an F_1_ population of 170 individuals derived from mixed intergeneric crosses between two sweet oranges (*Citrus sinensis* ‘Sanford’ and ‘Succari’) and two trifoliate oranges (*Poncirus trifoliata* ‘Argentina’ and ‘Flying Dragon’), of which 86 individuals were randomly chosen as a phenotyping population. All progenies were clonally propagated by grafting on ‘Volkamer’ lemon (*Citrus volkameriana*) in the greenhouse of USDA/ARS in Fort Pierce in 2010. In addition to citranges on ‘Volkamer’, ‘Volkamer’ lemon seedings, two sweet orange varieties (‘Navel’ grafted on ‘Swingle’ and ‘Hamlin’ grafted on C-35), and six trifoliate orange varieties (‘Flying-Dragon’ grafted on Volkamer, ‘Argentina’ grafted on Volkamer, ‘Pomeroy’ grafted on Volkamer, ‘Rubidoux’ seedlings, ‘Rich 16-6’ seedlings, and ‘Large-flower’ seedlings) were included. A completely randomized experiment of 768 trees with eight clonal replicate trees for each of the individuals and control varieties (‘Volkamer’ lemon seedings had sixteen clonal replicate trees) was established in the field trial of USDA-ARS in Fort Pierce in 2011. The plantation consisted of eight rows oriented south-north, with 4 m × 1.5 m planting distances. Guard trees not analyzed in this study were planted at the ends of each row. The field population trees were irrigated and fertilized with professional practices, but pesticides were not applied during the study, to encourage psyllid population increase, feeding, colonization, and inoculation of *C*Las to the trees. Maintenance of the field trees has been described elsewhere (Richardson et al. 2011).

### Evaluation of disease

The HLB disease pressure was high in the field trial of USDA-ARS in Fort Pierce, as reported in other concurrent studies (Lewis-Rosenblum *et al.*, 2015; Ramadugu *et al.*, 2016). The *C*Las-positive psyllid populations were adequate for naturally inoculating all the field materials with homogeneous disease pressure, providing excellent conditions to evaluate trees under natural disease spread conditions (Richardson et al. 2011, Westbrook et al. 2011). Beginning in early October 2013, disease evaluation was performed three times a year at intervals of 10 weeks, in late May, middle August and early October, representing the seasonal range in HLB symptoms and pathogen titers. At each time point, at least four fully expanded mature leaves were randomly collected from different branches and different quadrats of each tree for disease evaluation. The midribs were separated from the leaves and cut into small piece to extract DNA using the CTAB extraction method. The TaqMan label-based multiplex real-time PCR method was used to detect *C*Las titer in citrus leaves (Li *et al.*, 2006b; Li *et al.*, 2008). Real-time qPCR was perform on an ABI 7500 thermocycler with probes specific to *C*Las 16S ribosomal gene and citrus cytochrome oxidase gene. The mean cycle threshold (Ct) values of qPCR for direct *C*Las detection were normalized with Ct values of corresponding host plant gene. The Ct value of *C*Las detection, associated with titer of *C*Las in leaves, was employed as a phenotypic trait to evaluate the status of *C*Las infection. Trees with Ct value above 33 were considered to be HLB-negative (healthy tree) (Albrecht and Bowman, 2012). The percentage of healthy trees among eight replicate trees was employed as another phenotypic trait.

### Genotyping by sequencing

Genomic DNA of each citrange hybrid and parent was extracted using a modified CTAB method (Aldrich and Cullis, 1993), Genomic DNA was digested with restriction endonuclease and then processed into Restriction site Associated DNA (RAD) libraries according to a previously described protocol (Baird *et al.*, 2008). The constructed RAD libraries were sequenced on an Illumina HiSeq2000 platform following the manufacturer’s protocol. Data analysis and bioinformatics pipelines were provided by Floragenex (Floragenex Inc. Portland, Oregon, USA). Genotypes at each locus were determined using the VCF Popgen Pipeline version 4.0 to generate a customized VCF 4.1 (variant call format) database with parameters set as follows: minimum allele frequency for genotyping 0.075, minimum Phred score 15, minimum depth of sequencing coverage of 12, and minimum percentage of individuals with genotype calls 75%.

### Linkage analysis and construction of parental linkage maps

As extremely low genetic diversity was found between the different sweet orange parents and between the trifoliate orange parents, all F_1_ progenies from different crosses between the two genera were considered as a single family (Ollitrault *et al.*, 2012b). SNP markers segregating from only one of the parents were selected to construct the linkage maps. SNP markers matching the following criteria were excluded from linkage analysis: (1) had a missing genotype in more than 10% of progenies; (2) had a missing genotype for one of the parents; (3) had inconsistent genotypes among different parental sweet oranges or trifoliate oranges; (4) had homozygous genotypes for both parents; (5) had heterozygous genotype for both parents; (6) had unexpected segregation genotypes in more than 5% of progenies; (7) had no segregation in more than 95% of progenies. Markers with acceptable skewed segregation were included in linkage analysis referring to previous studies in citrus (Bernet *et al.*, 2010; Ollitrault *et al.*, 2012a; Raga *et al.*, 2012). Linkage analyses were performed using JoinMap 4.1 (Van Ooijen, 2011). Linkage mapping was performed under a two-way pseudo-testcross scheme (Grattapaglia and Sederoff, 1994) with two separate datasets, one with markers segregating from the trifoliate orange and the other one with markers segregating from sweet orange. Linked markers were grouped using the independence logarithm of the odds (LOD) with a threshold LOD score of 4.0 and a maximum recombination fraction (θ) of 0.4. Markers with identical segregation patterns or segregating similarity higher than 97% were excluded from the linkage groups. Map distances were estimated in centiMorgans (cM) using the regression mapping algorithm and the Kosambi mapping function. The linkage groups were numbered according to the corresponding scaffold number of the reference Clementine mandarin genome and marker names were composed of the scaffold number and SNP position on this reference. The genetic map was drawn using the MapChart 2.2 program (Voorrips, 2002)

### QTL mapping

QTL mapping was carried out on two parental maps separately using MapQTL 6 under the backcross model by composite interval mapping (IM) (van Ooijen, 1992). Phenotypic data were analyzed separately for each trait and each time point and year. The LOD thresholds to declare a significant QTL for each trait were determined via permutation tests using 1000 permutations at a genome-wide significance level of 0.90. The composite interval mapping analysis was performed with a step size of 1 cM to detect significant QTLs with a LOD score higher than the threshold. The nearest marker to the likelihood peak of each significant QTL was selected as a cofactor to perform restricted multiple QTL mapping (rMQM). If the LOD value linked with a cofactor fell below threshold during rMQM mapping, the cofactor was removed and the analysis repeated. This process was continued until the cofactor list remained stable. Kruskal-Wallis (KW) test was also used to provide complementary validation for significant genotypic means. Each significant QTL was characterized by its LOD score, percentage of explained phenotypic variation, confidence interval (in cM) corresponding to threshold LOD score and extension region at either side of the likelihood peak until the LOD score dropped to 2.0. QTLs with a LOD scores higher than 2.0 but not statistically significant were declared as minor QTLs. QTLs that showed clearly overlapping confidence intervals, close LOD peaks and similar allelic effects, were considered as co-localized.

### Data analysis

Descriptive statistics of all phenotypic data were calculated using SPSS. Value of phenotypic trait for each genotype at each time point was the averaged value of all replicate trees. Mean value for each year was the means of three time points within corresponding year. Max value for each year was the lowest Ct value (highest CLas titer) of three time points within corresponding year. To evaluate whether the data followed a normal distribution, a normality analysis by Kolmogorov-Smirnov and Shapiro-Wilk tests was performed separately for each of the traits at each of the time points and years. Histograms for each trait were constructed using the complete data set. Pearson’s correlation coefficients were calculated between each of the time points and years vs. the two phenotypic traits. The Box-Cox transformation was performed before QTL analysis if data presented a non-normal distribution.

## Results

### RAD sequencing and SNP-based genotyping

Four parental varieties (‘Argentina’ and ‘Flying Dragon’ trifoliate orange; ‘Sanford’ and ‘Succari’ sweet orange) and 188 putative hybrids were processed for RAD sequencing via NGS Illumina platform. Sequencing data indicated that 18 of the putative hybrids were triploid hybrids, multiple-cross hybrids, self-pollinated, or clonal individuals. These individuals were eliminated from further analysis, resulting in a total of 170 true F_1_ progenies from the cross of trifoliate orange and sweet orange. Except for one progeny with poor quality of sequencing reads, all had acceptable quality of sequencing data. The average number of reads was 8.14E6 and 7.37E6 for trifoliate orange and sweet orange respectively. The read counts for 169 F_1_ progenies ranged from 1.05E6 to 14.91E6, with an average of 4.53E6 reads. The average number of clusters was 1.83E5 and 1.47E5 for trifoliate orange and sweet orange respectively, and the cluster numbers for the F_1_ progenies ranged from 0.32E5 to 2.79E5 and averaged to be 1.03E5.

SNP calling of parents and progenies was performed with standard filtering criteria using the Clementine mandarin genome as a reference. In total, 62,711 putative SNP loci were determined, of which 55.6% were transitions and 44.4% were transversions. Of the putative loci, 96.6% were identical between ‘Argentina’ and ‘Flying Dragon’ trifoliate orange, and 98.0% were identical between ‘Sanford’ and ‘Succari’ sweet orange. Considering low genetic diversity were found within different trifoliate orange parents and sweet orange parents, all the F_1_ progenies from different crosses between trifoliate orange and sweet orange were considered as a single family. The marker configuration codes ‘lm×ll’ and ‘nn×np’ were used to represent markers that were heterozygous only in one of parents. SNP markers heterozygous in both parents (with the configuration code ‘hk×hk’) were not considered in this study to ensure high quality of genotyping and linkage mapping for each parent. After stringent filtering, a total of 3861 SNP markers with configuration code ‘lm×ll’ or ‘nn×np’ were retained for genetic map construction, of which 1408 SNP markers were heterozygous only in trifoliate orange and 2453 SNP markers were heterozygous only in sweet orange.

### Genetic linkage map construction and evaluation

After eliminating markers with identical or similar segregation patterns, as well as markers with weak linkage, SNP markers with high quality in each dataset were grouped into nine linkage groups under the threshold LOD score of 4.0, which is consistent with the number of chromosomes in citrus. Additionally, all nine linkage groups conserved their integrity up to LOD of 10 for both parents. Finally, for trifoliate orange, a total of 647 high-quality SNP markers were mapped on nine linkage groups with unique loci, spanning a total genetic length of 1030.8 cM with an average inter-locus distance of 1.59 cM (Table 1 and Fig. 1). The number of markers within each linkage group ranged from 52 (for LG-t6) to 125 (for LG-t3), spanning a genetic distance ranging from 78.7 cM (for LG-t6) to 155.2 cM (for LG-t3). In total, 85.5% of the inter-loci gaps on the genome-wide genetic map were smaller than 3 cM and no gap was larger than 10 cM. For sweet orange, 754 high-quality SNP markers with unique loci were mapped into nine linkage groups, spanning a total genetic length of 760.2 cM with an average inter-loci distance of 1.01 cM (Table 1 and Fig. 2). The number of markers within each linkage group ranged from 57 (for LG-s1) to 118 (for LG-s3), spanning a genetic distance ranging from 60.5 cM (for LG-s9) to 105.3 cM (for LG-s3). Of the inter-locus gaps on the whole genetic map, 93.3% were smaller than 3 cM and only one gap was larger than 10 cM (11.2 cM on LG-s1).

**Table 1.** Summary of the genetic linkage maps of trifoliate orange and sweet orange.

| Linkage Group |  | Reference genome | Number of marker loci |  | LG Length (cM) |  | Average inter-loci distance (cM) |  | Number of Syntenic marker |  |
| --- | --- | --- | --- | --- | --- | --- | --- | --- | --- | --- |
| Trifoliate orange | Sweet orange | Clementine mandarin | Trifoliate orange | Sweet orange | Trifoliate orange | Sweet orange | Trifoliate orange | Sweet orange | Trifoliate orange | Sweet orange |
| LG-t1 | LG-s1 | Scaffold_1 | 70 | 57 | 117.9 | 93.4 | 1.68 | 1.64 | 70 | 57 |
| LG-t2 | LG-s2 | Scaffold_2 | 87 | 89 | 124.5 | 99.6 | 1.43 | 1.12 | 87 | 89 |
| LG-t3 | LG-s3 | Scaffold_3 | 125 | 118 | 155.2 | 105.3 | 1.24 | 0.89 | 125 | 118 |
| LG-t4 | LG-s4 | Scaffold_4 | 64 | 94 | 121.5 | 74.0 | 1.90 | 0.79 | 64 | 88 |
| LG-t5 | LG-s5 | Scaffold_5 | 72 | 98 | 123.9 | 90.9 | 1.72 | 0.93 | 72 | 98 |
| LG-t6 | LG-s6 | Scaffold_6 | 52 | 77 | 78.7 | 68.2 | 1.51 | 0.89 | 52 | 77 |
| LG-t7 | LG-s7 | Scaffold_7 | 57 | 85 | 108.3 | 86.2 | 1.90 | 1.01 | 47 | 73 |
| LG-t8 | LG-s8 | Scaffold_8 | 58 | 76 | 102.8 | 82.1 | 1.77 | 1.08 | 58 | 64 |
| LG-t9 | LG-s9 | Scaffold_9 | 62 | 60 | 98.0 | 60.5 | 1.58 | 1.01 | 62 | 60 |
| Total |  |  | 647 | 754 | 1030.8 | 760.2 | 1.59 | 1.01 | 637 | 724 |

The correspondence between linkage groups and syntenic scaffolds of Clementine mandarin genome was based on the location of most mapped markers.

**Fig 1.**
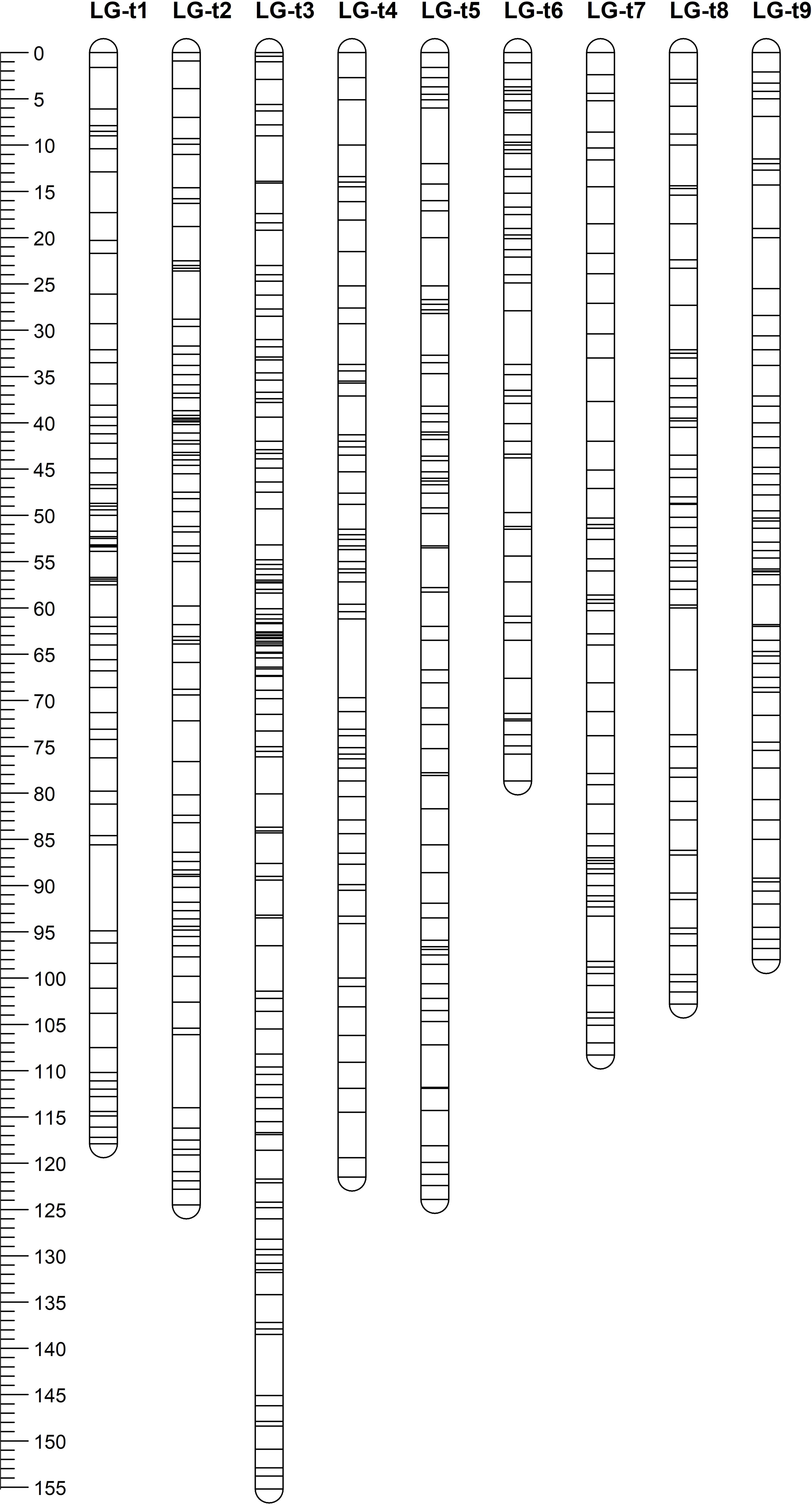
Distribution of markers with unique loci in the trifoliate orange genetic linkage map. The nine linkage groups correspond to the nine major scaffolds of the Clementine mandarin genome. Map distances in centiMorgans (cM).are indicated by the ruler at the left.

**Fig 2.**
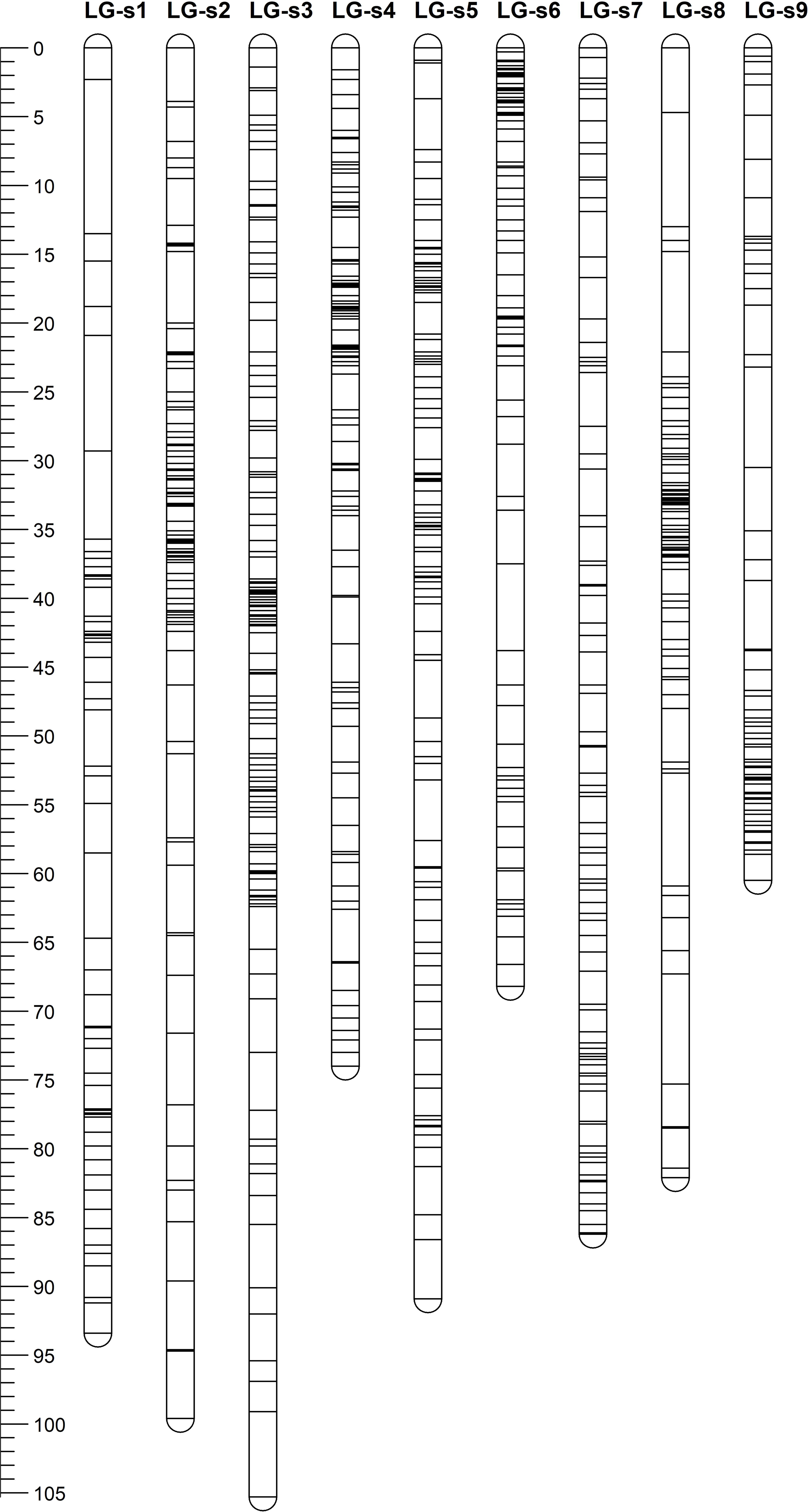
Distribution of markers with unique loci in the sweet orange genetic linkage map. The nine linkage groups correspond to the nine major scaffolds of the Clementine mandarin genome. Map distances in centiMorgans (cM).are indicated by the ruler at the left.

Through alignment, all SNP markers on the two genetic maps were successfully mapped onto nine major scaffolds of the Clementine mandarin genome. For trifoliate orange, except for 10 markers of LG-t7 which mapped on Scaffold_5, all other 637 markers were mapped onto syntenic scaffolds (Table 1 and Fig. 3). The overall coverage ratio of mapped markers on the Clementine genome was 98.6%, and the coverage ratio on each scaffold ranged from 95.5% (for LG-t7) to 99.7% (for LG-t5). For sweet orange, except for a total of 30 markers on LG-s4, LG-s7 and LG-s8, all other 724 markers were mapped onto syntenic scaffolds (Table 1 and Fig. 4). The overall coverage ratio of mapped markers on the Clementine genome was 95.0%, and the coverage ratio on each scaffold ranged from 87.7% (for LG-s9) to 99.3% (for LG-s8). Only some minor discrepancies were observed between the genetic maps and the Clementine genome, indicating possible genetic diversity among different citrus species, or potential errors in the current linkage grouping, or erroneous assemblies in Clementine mandarin genome. Collinear analysis of the consensus between the genetic map and the Clementine genome via dot-plot diagram not only showed variations of the ratios between genetic distance to physical distance, but also clearly revealed high conservation of synteny between the genetic maps of trifoliate orange and sweet orange.

**Fig 3.**
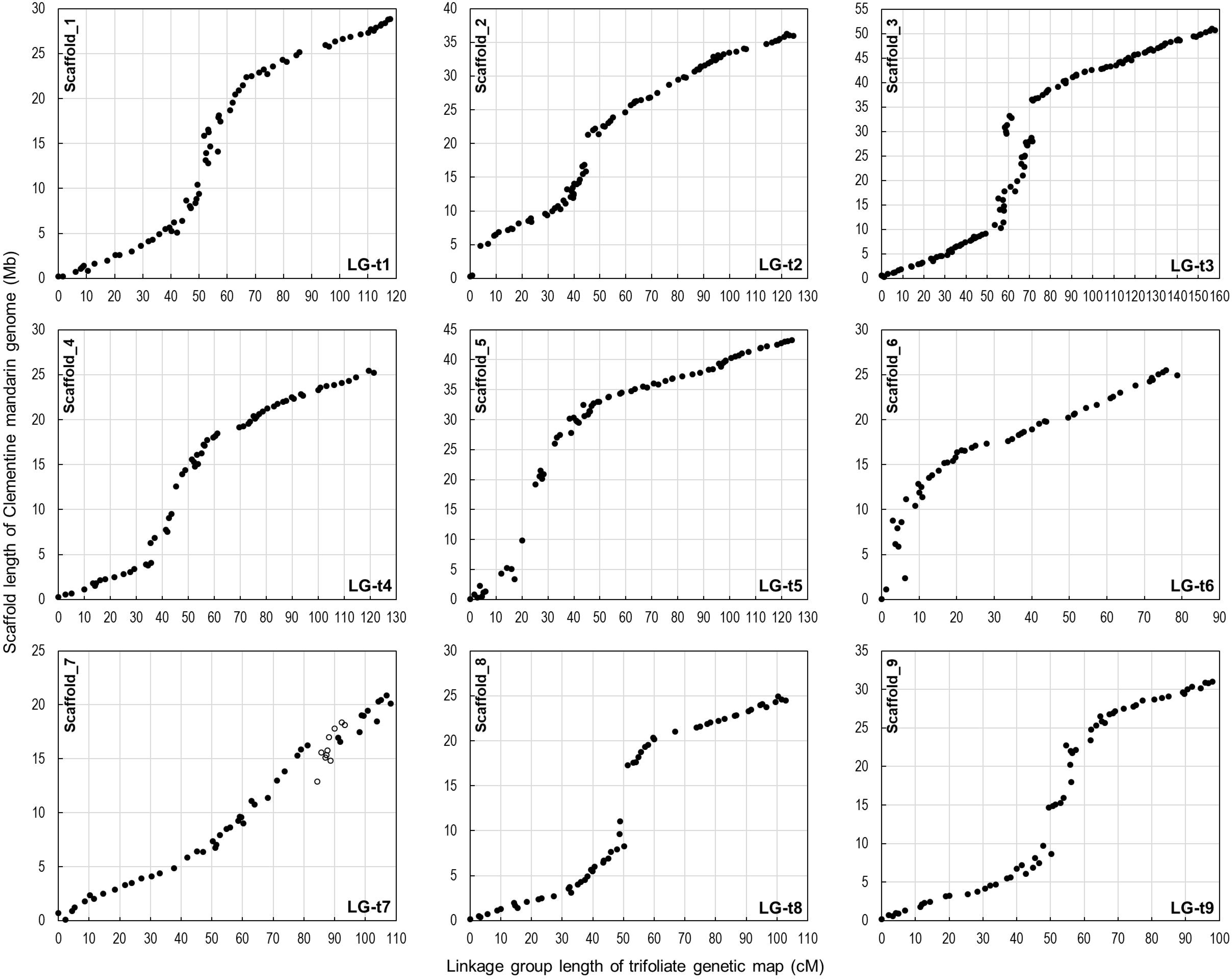
Collinear analysis of the consensus between the trifoliate orange genetic linkage map and the Clementine mandarin genome via dot-plot diagram. Solid spots represent markers that mapped on syntenic scaffolds, while hollow spots represent markers that mapped on other scaffolds (LG7 versus Scaffold_5).

**Fig 4.**
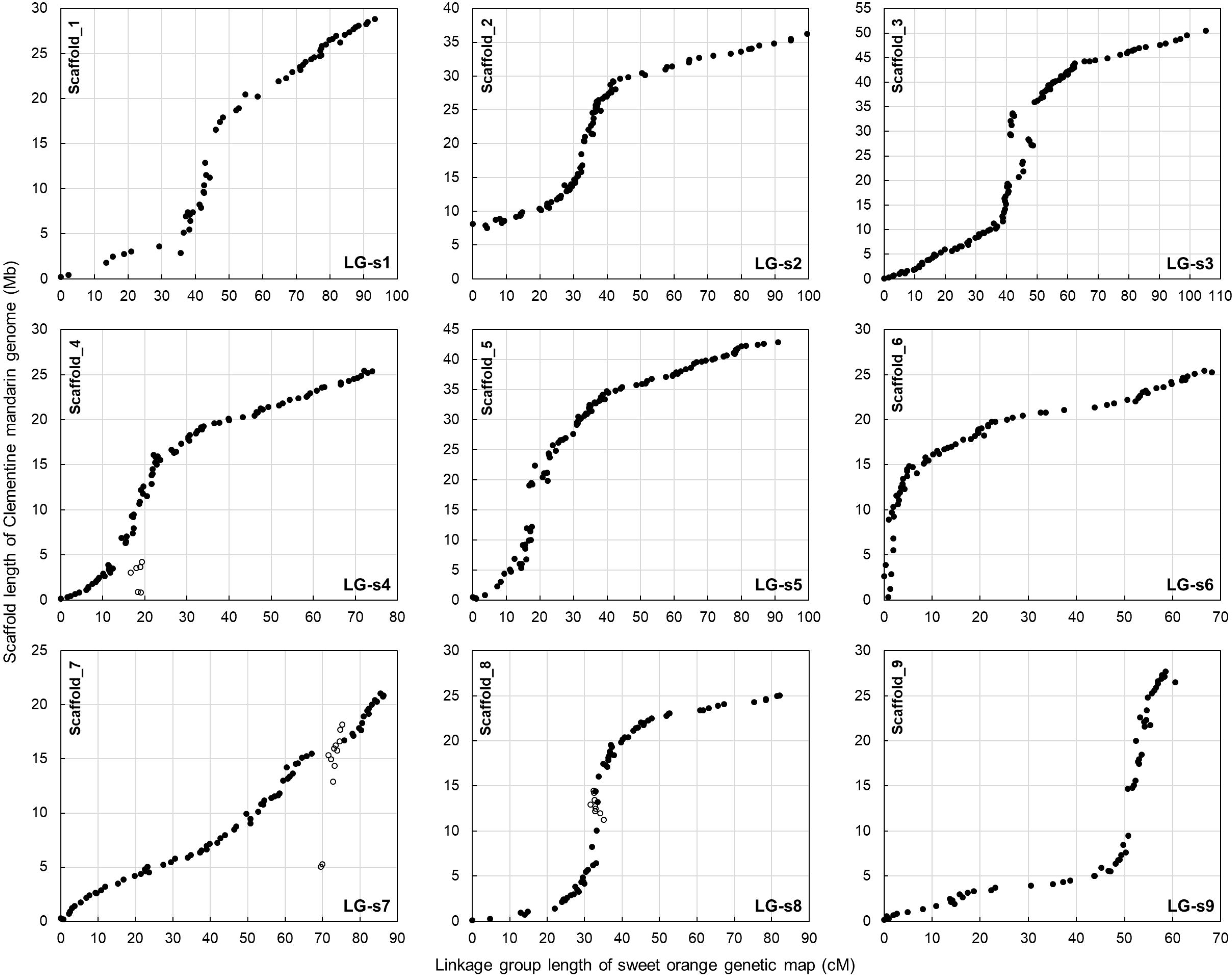
Collinear analysis of the consensus between the sweet orange genetic linkage map and the Clementine mandarin genome via dot-plot diagram. Solid spots represent markers that mapped on syntenic scaffolds, while hollow spots represent markers that mapped on other scaffolds.

### Phenotyping of *C* Las infection

The phenotyping population contained a random collection of 86 F_1_ progenies from the mapping population, as well as six trifoliate oranges, two sweet oranges, and the rootstock seedings, with eight clonal replicate trees for each of individuals. After exposure to intense HLB pressure for two to five years in a replicated field trial, the phenotyping population was evaluated for *C*Las infection in leaves by qPCR from October 2013 to October 2016, three times per year and ten times in total. Table 2 summarizes Ct values associated with *C*Las titer in leaves at different time points and years. Throughout the evaluation period, all sweet orange varieties were severely infected by *C*Las with Ct values under 28 constantly and averaging 25.7, while all trifoliate orange varieties were HLB negative with Ct values above 35 constantly and averaging 37.6. For F_1_ progenies, Ct values ranged from 24 to 38, and the average Ct value of all progenies fluctuated between 32 and 27 with higher phenotypic variation found at earlier time points than later time points. The frequency distribution of Ct value among the F_1_ progenies at each of the time points and years are illustrated in Fig. 5. As shown, obvious segregation for Ct values was observed for all time points, especially at earlier time points.

**Table 2.** Ct value of *C*Las detection in control varieties and F_1_ progenies at different time points and years.

|  | **Hamlin<sup>a</sup>** | **Navel<sup>a</sup>** | **Argentina<sup>a</sup>** | **Flying-Dragon<sup>a</sup>** | **Large-flower<sup>a</sup>** | **Pomeroy<sup>a</sup>** | **Rich 16-6<sup>a</sup>** | **Rubidoux<sup>a</sup>** | **Volkamer<sup>a</sup>** | **Progenies<sup>b</sup>** |
| --- | --- | --- | --- | --- | --- | --- | --- | --- | --- | --- |
|  | Sweet orange | Sweet orange | Trifoliolate orange | Trifoliolate orange | Trifoliolate orange | Trifoliolate orange | Trifoliolate orange | Trifoliolate orange | Rootstock | Hybrids |
| Ct-2013-Oct | 26.2 ± 5.6 | 23.3 ± 0.9 | 39.8 ± 0.7 | 38.9 ± 3.0 | 40.0 ± 0.0 | 39.8 ± 0.6 | 39.6 ± 0.8 | 39.2 ± 2.3 | 27.4 ± 5.5 | 32.0 ± 7.1 |
| Ct-2014-May | 27.0 ± 6.4 | 27.7 ± 6.6 | 38.0 ± 2.2 | 38.2 ± 2.9 | 37.4 ± 2.5 | 37.9 ± 2.4 | 36.1 ± 5.5 | 37.0 ± 3.5 | 30.2 ± 5.8 | 30.6 ± 6.6 |
| Ct-2014-Aug | 26.6 ± 4.4 | 24.3 ± 4.0 | 40.0 ± 0.0 | 38.6 ± 1.9 | 37.8 ± 2.5 | 38.6 ± 2.9 | 38.9 ± 2.0 | 38.7 ± 2.5 | 31.0 ± 5.1 | 29.1 ± 6.3 |
| Ct-2014-Oct | 25.8 ± 0.7 | 25.1 ± 2.8 | 39.4 ± 1.8 | 39.4 ± 1.5 | 38.4 ± 3.7 | 36.9 ± 5.3 | 38.1 ± 4.5 | 38.2 ± 1.6 | 27.4 ± 3.0 | 28.1 ± 5.9 |
| Ct-2015-May | 25.3 ± 2.1 | 23.0 ± 1.6 | 36.8 ± 5.8 | 36.8 ± 5.3 | 37.2 ± 6.0 | 35.7 ± 5.9 | 35.4 ± 6.0 | 36.6 ± 6.2 | 28.6 ± 4.5 | 28.4 ± 5.5 |
| Ct-2015-Aug | 26.3 ± 2.1 | 26.4 ± 5.4 | 37.2 ± 4.8 | 38.0 ± 3.4 | 39.6 ± 1.1 | 37.8 ± 4.5 | 38.3 ± 4.0 | 36.0 ± 5.8 | 29.5 ± 5.2 | 30.2 ± 6.1 |
| Ct-2015-Oct | 25.0 ± 2.3 | 24.6 ± 2.1 | 35.7 ± 5.1 | 39.8 ± 0.6 | 37.2 ± 5.2 | 37.4 ± 4.9 | 37.8 ± 5.2 | 35.6 ± 5.7 | 28.1 ± 2.7 | 28.0 ± 4.9 |
| Ct-2016-May | 27.2 ± 1.3 | 27.4 ± 1.8 | 40.0 ± 0.0 | 38.2 ± 4.7 | 36.9 ± 5.4 | 37.2 ± 5.2 | 38.3 ± 4.8 | 37.3 ± 5.0 | 31.7 ± 5.4 | 28.5 ± 3.7 |
| Ct-2016-Aug | 27.3 ± 1.5 | 26.8 ± 1.5 | 35.9 ± 5.7 | 38.9 ± 3.0 | 35.4 ± 5.2 | 34.4 ± 5.1 | 36.8 ± 5.3 | 35.1 ± 3.3 | 30.3 ± 3.8 | 29.0 ± 3.4 |
| Ct-2016-Oct | 24.0 ± 0.6 | 24.8 ± 1.0 | 35.5 ± 5.6 | 36.8 ± 3.8 | 37.2 ± 4.8 | 33.4 ± 5.6 | 36.5 ± 3.7 | 38.0 ± 2.0 | 27.3 ± 2.8 | 27.4 ± 3.3 |
| Ct-2014-Mean<sup>c</sup> | 26.5 ± 4.8 | 25.4 ± 4.3 | 39.1 ± 1.1 | 38.7 ± 1.3 | 37.9 ± 1.8 | 37.8 ± 2.0 | 37.7 ± 3.6 | 37.9 ± 1.2 | 29.5 ± 3.3 | 29.2 ± 4.6 |
| Ct-2015-Mean<sup>c</sup> | 25.2 ± 1.8 | 24.7 ± 1.4 | 36.4 ± 4.1 | 38.3 ± 2.8 | 37.6 ± 3.1 | 36.7 ± 3.3 | 36.5 ± 4.7 | 36.1 ± 5.0 | 29.3 ± 3.1 | 28.7 ± 4.0 |
| Ct-2016-Mean<sup>c</sup> | 26.4 ± 1.1 | 26.2 ± 1.2 | 37.8 ± 2.8 | 38.1 ± 3.7 | 36.5 ± 3.9 | 35.6 ± 4.9 | 37.2 ± 4.6 | 36.7 ± 4.3 | 29.8 ± 2.6 | 28.3 ± 2.9 |
| Ct-2014-16-Mean<sup>d</sup> | 26.2 ± 2.5 | 25.3 ± 2.7 | 37.9 ± 3.3 | 38.5 ± 2.6 | 37.0 ± 3.1 | 36.8 ± 3.6 | 37.1 ± 4.2 | 36.9 ± 4.0 | 29.3 ± 3.1 | 28.8 ± 3.7 |
| Ct-2014-Max<sup>e</sup> | 22.5 ± 2.4 | 22.5 ± 3.3 | 37.8 ± 2.4 | 37.0 ± 2.7 | 35.9 ± 3.5 | 35.5 ± 3.6 | 36.2 ± 3.9 | 35.4 ± 2.7 | 25.4 ± 1.9 | 25.5 ± 4.3 |
| Ct-2015-Max<sup>e</sup> | 22.7 ± 2.6 | 21.8 ± 1.9 | 34.6 ± 4.9 | 36.8 ± 4.3 | 35.7 ± 4.4 | 34.3 ± 5.3 | 34.4 ± 5.3 | 33.9 ± 6.1 | 27.4 ± 3.5 | 25.4 ± 3.3 |
| Ct-2016-Max<sup>e</sup> | 23.5 ± 2.5 | 22.9 ± 3.1 | 35.2 ± 4.8 | 36.9 ± 4.9 | 33.8 ± 5.3 | 32.6 ± 6.2 | 35.7 ± 4.5 | 34.1 ± 5.8 | 26.6 ± 1.8 | 26.0 ± 2.5 |
| Ct-2014-16-Max-M<sup>f</sup> | 22.9 ± 2.1 | 22.4 ± 2.6 | 35.9 ± 3.3 | 36.9 ± 2.9 | 35.1 ± 4.0 | 34.1 ± 2.6 | 35.4 ± 3.7 | 34.5 ± 4.7 | 26.7 ± 2.1 | 25.6 ± 3.0 |

<sup>a</sup> The Ct value for these control varieties, shown as Mean ± SD, is based on evaluation of all replicate trees
<sup>b</sup> The Ct value for F<sub>1</sub> progenies is the averaged Ct value of 86 individuals.
<sup>c</sup> The averaged Ct values of three time points within corresponding year
<sup>d</sup> The averaged Ct value of nine time points of the three years
<sup>e</sup> The lowest Ct value (highest CLas titer) of three time points within corresponding year
<sup>f</sup> The averaged lowest Ct value of the three years

**Fig 5.**
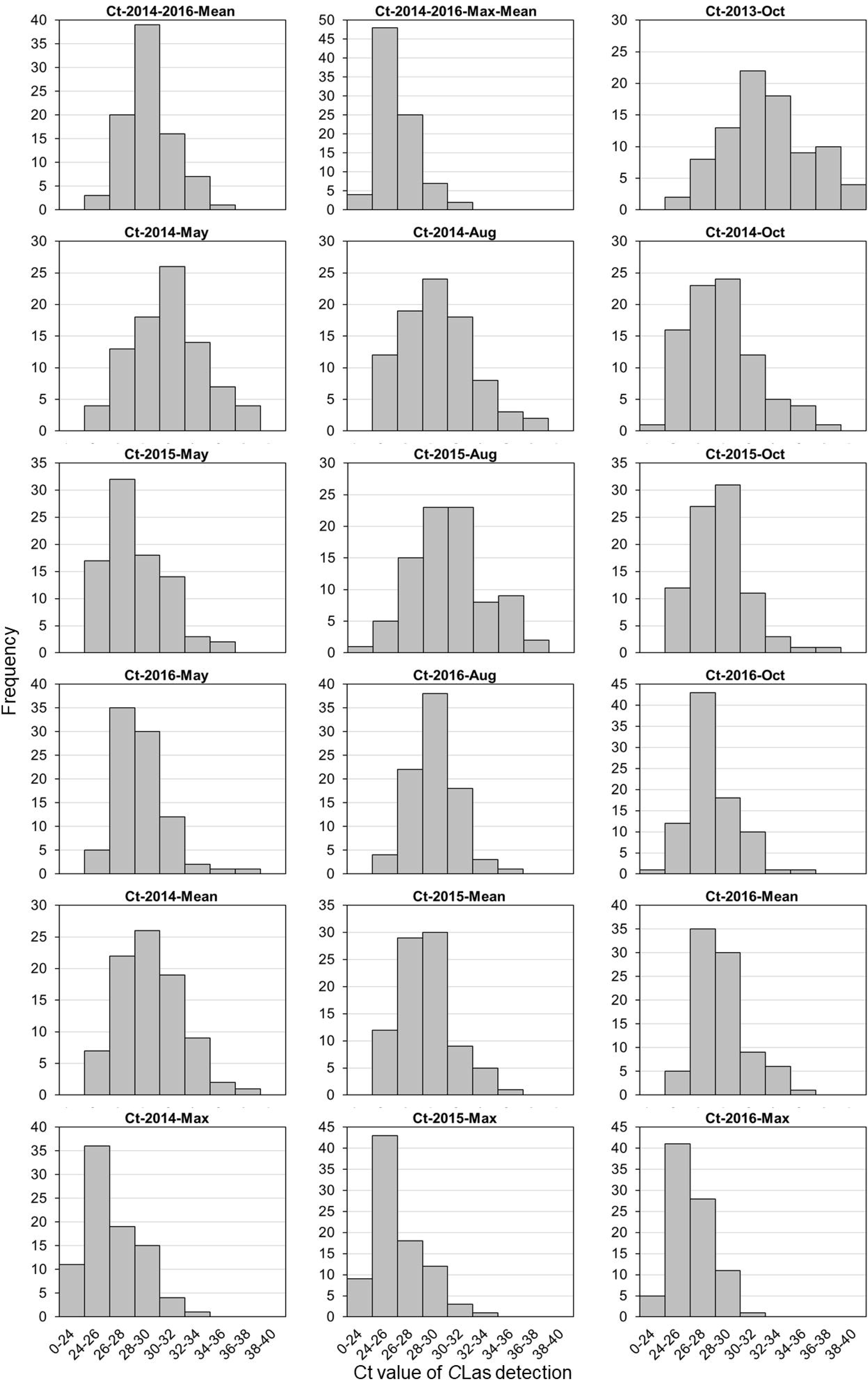
Frequency distributions of Ct value of *C*Las detection in F_1_ progenies at different time points and years.

Due to random factors associated with inoculation by psyllid vector, replicate trees of the same genotypes were not naturally inoculated at the same time. As a result, extremely high phenotypic variations were observed within replicate trees of the same genotypes. Thus, the percentage of healthy trees was employed as another phenotypic trait to reflect the efficiency and status of *C*Las infection. The percentage of healthy trees at each of the time points and years are summarized in Supplementary Table 1, and the frequency distribution of this trait among the F_1_ progenies is illustrated in Supplementary Fig. 1. As shown, throughout the evaluation period, the percentage of healthy trees was under 40% constantly and 12.9% in average for all sweet orange varieties, while it was above 60% constantly and 84.7% in average for all trifoliate orange varieties. For F_1_ progenies, the average mean percentage of healthy trees of all progenies fluctuated between 48% and 10% with higher phenotypic variation found at earlier time points.

Correlation coefficients of phenotypic data between time points and years were shown in Supplementary Table 2. Notably, very high correlation was found between the two traits at each of the time points and years with correlation coefficients all above 0.93. However, correlation among the ten time points was very low. The correlation coefficients ranged from 0.38 to 0.80 and 0.61 in average for the Ct value of *C*Las detection, and ranged from 0.36 to 0.73 and 0.59 in average for percentage of healthy trees, indicating high variation of seasonal *C*Las infection. The lowest correlations were mostly found with data of time point 2013-Oct and 2015-Aug. Correlation among the three years was relatively high. The average correlation coefficients were 0.79 and 0.82 respectively for the two traits, indicating overall stability of long-term *C*Las infection.

### Detection of QTLs associated with *C*Las infection

QTLs associated with *C*Las infection were detected separately in two parental genetic maps for each time point, year and phenotypic trait. In total, 14 significant QTLs were identified on the trifoliate orange genetic map (Table 3) and 16 significant QTLs were identified on the sweet orange genetic map (Table 4). In addition, as complementary support of QTL identification, 21 minor QTLs in trifoliate orange and 14 minor QTLs in sweet orange were detected, and most of them located in the same regions where significant QTLs exist. The map locations of 35 QTLs detected in trifoliate orange are shown in Fig. 6, while the map locations of 30 QTLs detected in sweet orange are shown in Fig. 7. Most of the QTLs for the two traits were co-localized. The maximum LOD score for each of the significant QTLs ranged from 2.6 to 6.3. The estimates of the phenotypic variation (R^2^) explained by each of the significant QTLs ranged from 13.7% to 32.8%. For the minor QTLs, the estimates of phenotypic variation explained ranged from 9.3% to 16.6%.

**Table 3.**
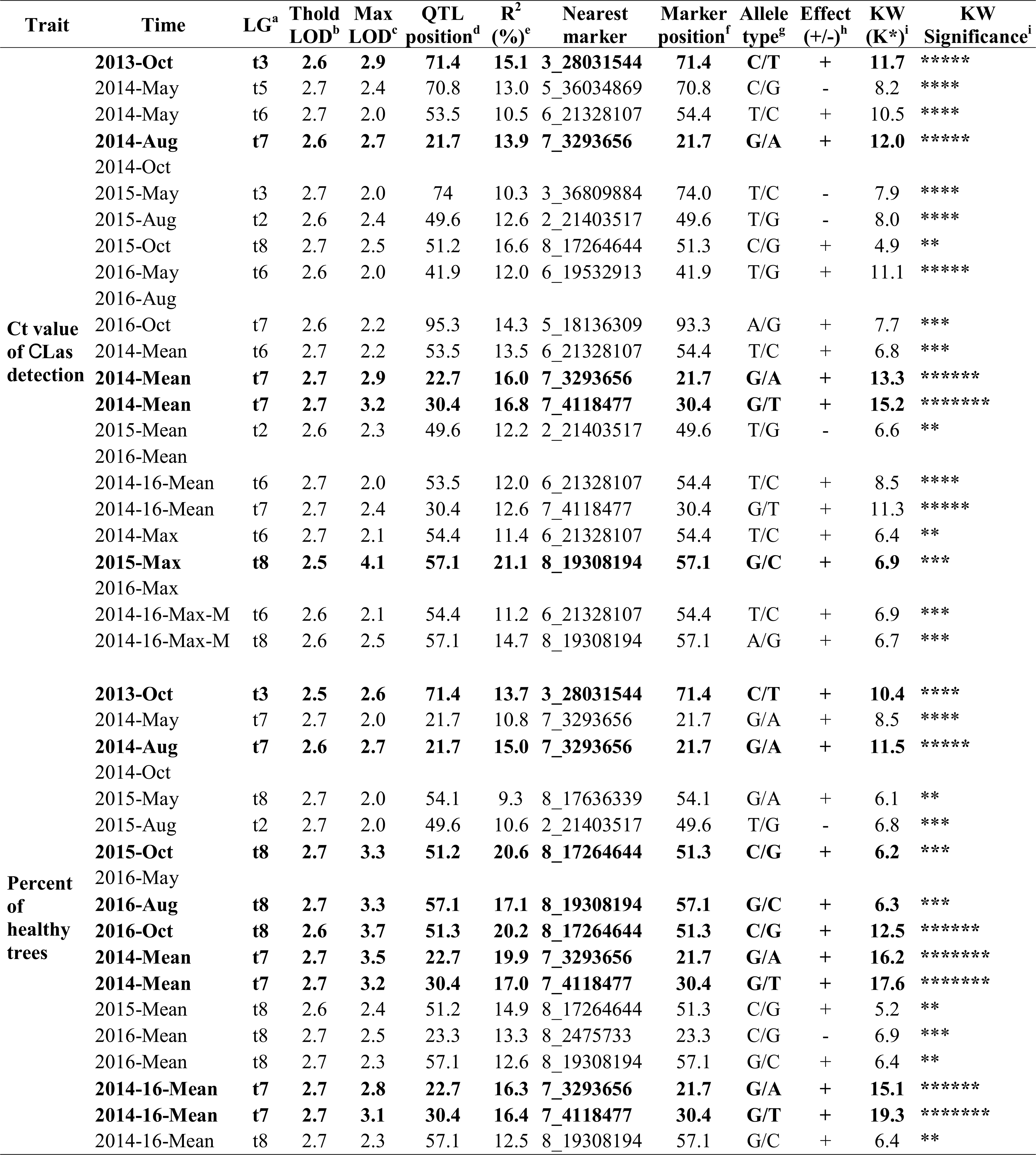
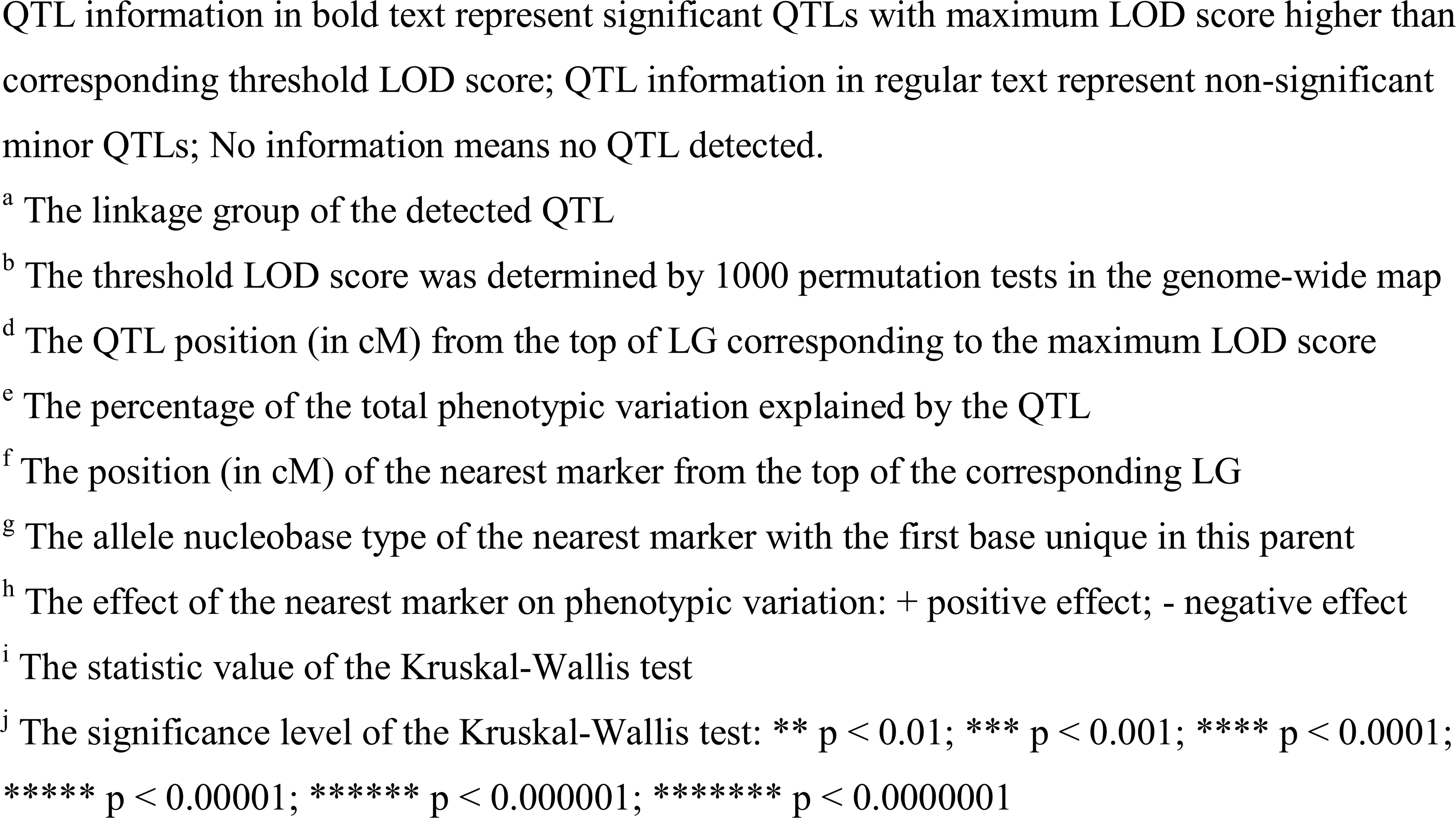
QTLs detected at different time points and years in trifoliate orange genetic map.

**Table 4.**
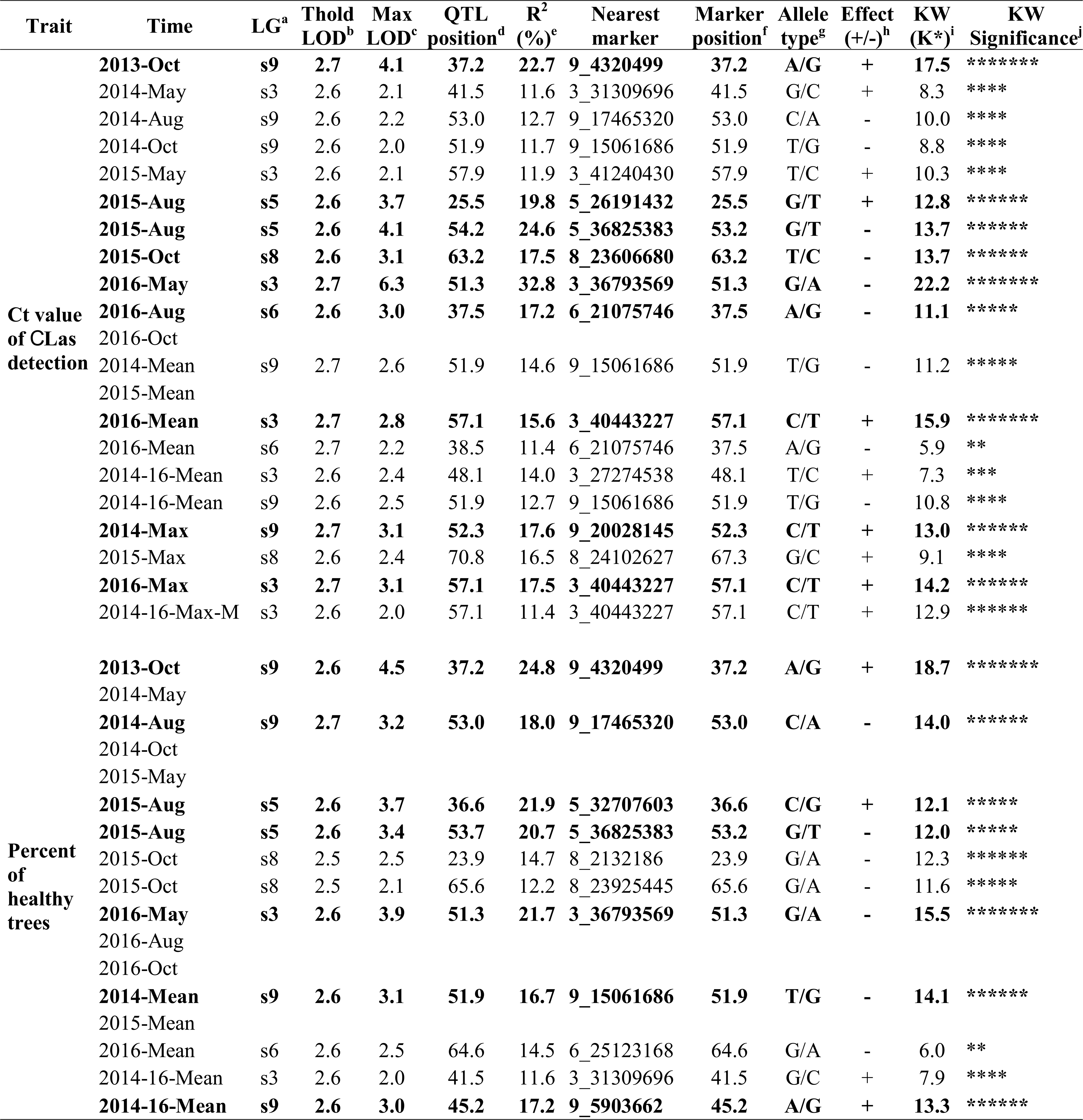
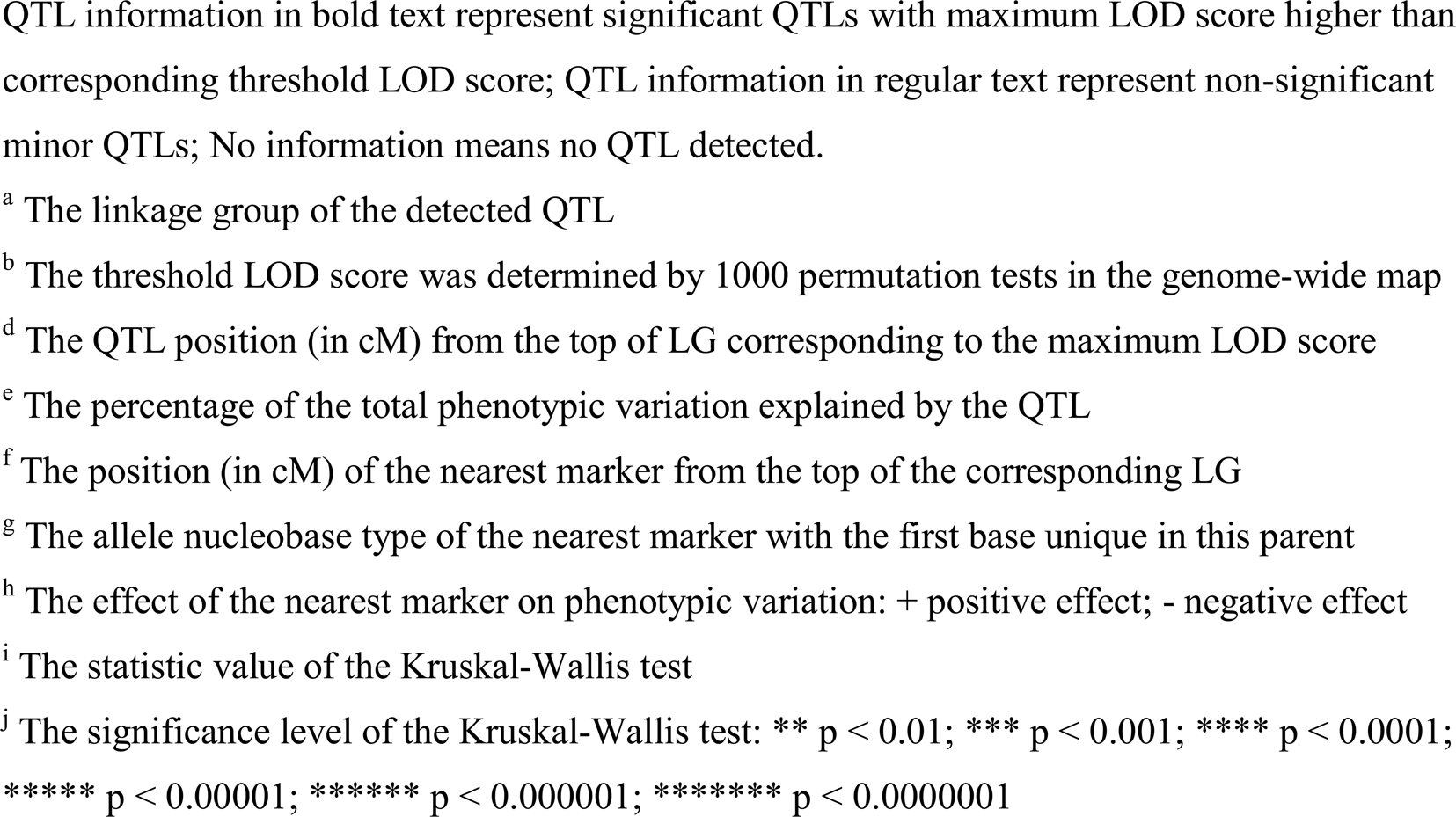
QTLs detected at different time points and years in sweet orange genetic map.

**Fig 6.**
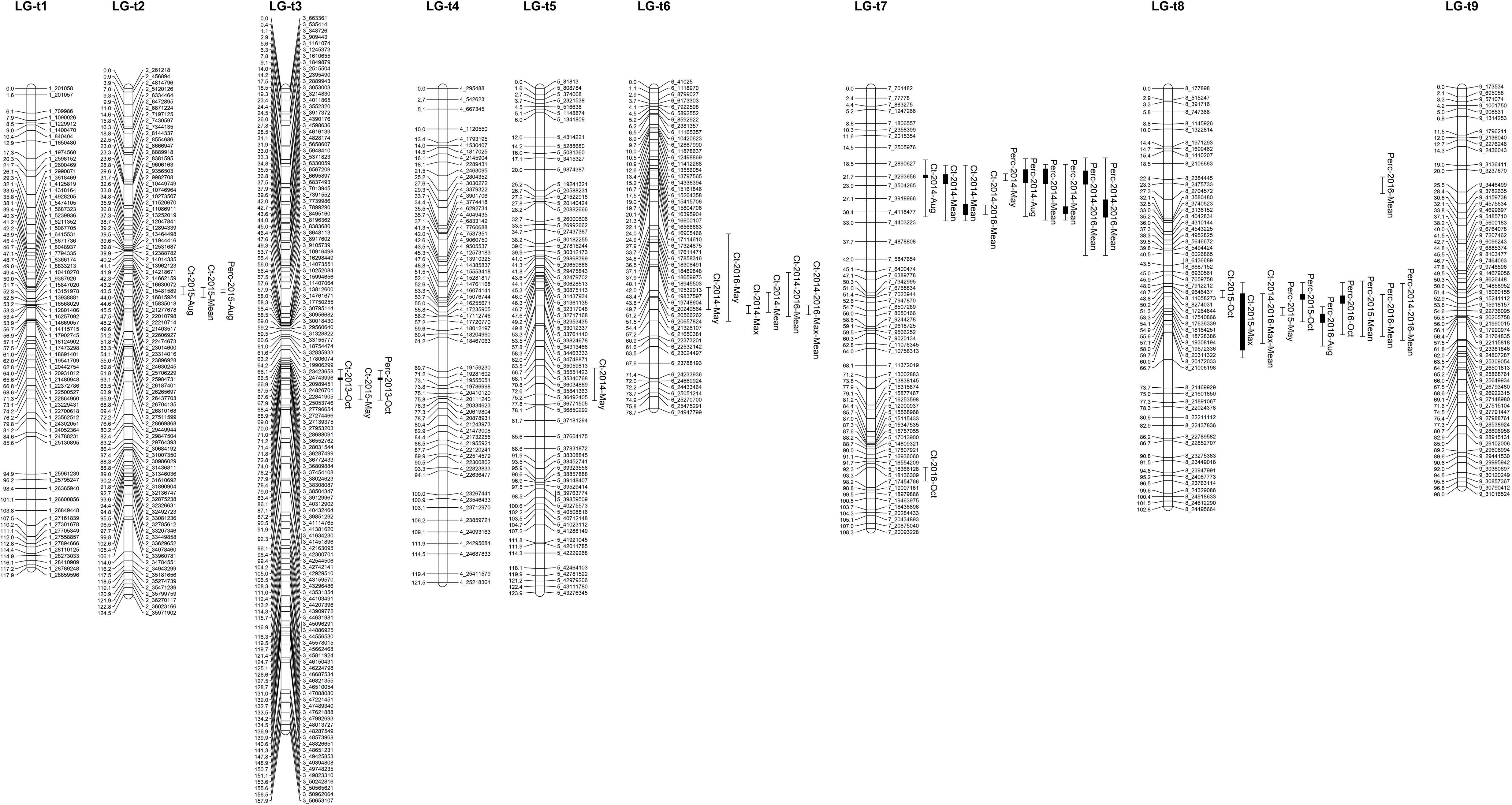
Mapping QTLs with Ct value of *C*Las detection and percentage of healthy trees on the trifoliate orange genetic linkage map. Thick bars on the right side of each LG indicate significant QTLs with LOD score above corresponding threshold and flanking error bars indicate extension of QTL region at LOD score 2.0. Thin single bars indicate minor QTLs with LOD scores between 2.0 and threshold.

**Fig 7.**
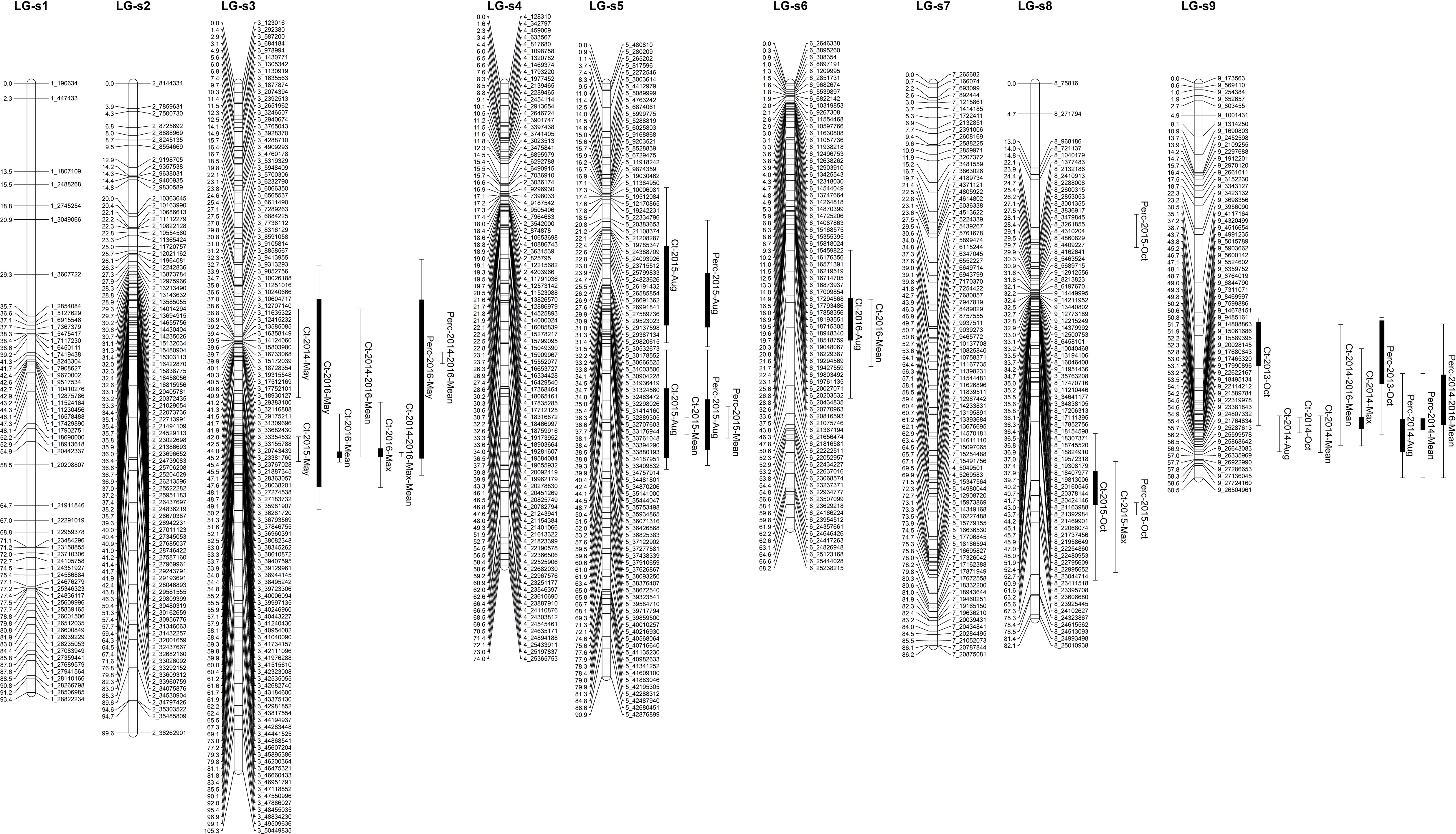
Mapping QTLs with Ct value of *C*Las detection and percentage of healthy trees on the sweet orange genetic linkage map. Thick bars on the right side of each LG indicate significant QTLs with LOD score above corresponding threshold and flanking error bars indicate extension of QTL region at LOD score 2.0. Thin single bars indicate minor QTLs with LOD scores between 2.0 and threshold.

Integrating the QTLs of two traits detected at different time points, all 14 significant QTLs on the trifoliate orange genetic map could be grouped into three main clusters, respectively located on LG-t3, LG-t7 and LG-t8; while the 16 significant QTLs on the sweet orange genetic map could be grouped into six clusters, respectively located on LG-s3, LG-s5, LG-s6, LG-s8 and LG-s9. In addition, most of the minor QTLs were also located in these clusters. Co-localization of QTLs between two traits were observed in all three clusters in trifoliate orange and four of the six clusters in sweet orange (located on LG-s3, LG-s5 and LG-s9). Of the nine QTL clusters, five QTL clusters, respectively located on LG-t7 and LG-t8 of the trifoliate orange genetic map and LG-s3, LG-s5 and LG-s9 of the sweet orange genetic map, collectively explained a major part of the phenotypic variation. However, repeatability of QTLs among three years was quite limited for either of the traits. The best repeatable QTL cluster was located on LG-t8 with three significant QTLs for the percentage of healthy trees detected in two years.

Further examining these significant QTLs, in trifoliate orange the five nearest markers (3_28031544, 7_3293656, 7_4118477, 8_17264644 and 8_19308194) all exhibited positive effect on the phenotypic traits (Table 3). While in sweet orange, the situation is more complex; among the 11 nearest markers (3_36793569, 3_40443227, 5_26191432, 5_32707603, 5_36825383, 6_21075746, 8_23606680, 9_4320499, 9_5903662, 9_15061686 and 9_17465320), six exhibited negative effects on the phenotypic traits and five exhibited positive effects (Table 4). Notably, all of these nearest markers were significantly associated with corresponding phenotypic data under the Kruskal-Wallis test. The above results indicate that both trifoliate orange and sweet orange genetically contributed to the tolerance of the progenies to *C*Las infection.

## Discussion

### Genotyping and linkage mapping

In this study, we constructed two separate parental genetic maps, which we believe to be the highest density genetic maps to date for trifoliate orange and sweet orange. The former highest density genetic map of trifoliate orange consisted of 146 SNP markers and 74 SSR markers spanning a total genetic length of 937.1 cM with an average density of 4.26 marker/cM (Lyon, 2008). The former highest density genetic map of sweet orange consisted of 799 SNP markers and 189 SSR markers at 972 loci spanning a total genetic length of 1026.6 cM with an average density of 1.06 marker/cM (Lyon, 2008), which was used as a reference map for the assembly of a sweet orange genome (Wu *et al.*, 2014; Xu *et al.*, 2013). The saturation of trifoliate orange genetic map is greatly improved in our study, which could be utilized as a reference map for assembling the trifoliate orange genome in the future. However, citrus genetic maps require substantial additional effort to achieve the high-density and high-resolution found in model and agronomic species genetic maps, that include thousands of markers with high accuracy and precision. Although we initially developed numerous SNP markers for our linkage mapping, only a small portion of markers were successfully mapped as unique loci. This has been observed in all previous SNP-based genetic maps of citrus, where many markers were located in ‘zero recombination clusters’, indicating nearly no meiotic recombination occurs between markers within those regions (Curtolo *et al.*, 2017; Guo *et al.*, 2015; Imai *et al.*, 2017; Lyon, 2008; Shimada *et al.*, 2014; Yu *et al.*, 2016). High redundancy of identified markers is commonly attributed to both small population size and closely adjacent physical marker location, and could be significantly improved by increasing the size of mapping populations and the evenness of marker distribution. These solutions have been applied to mapping of many plant species, but are difficult for citrus species. Guo et al successfully constructed an integrated genetic map of pummelo with 1543 SNP markers using an intraspecific full-sib F_1_ population with only 124 individuals (Guo *et al.*, 2015). By contrast, Curtolo et al. attempted to construct a genetic map using an interspecific full-sib F_1_ population with 278 individuals from crossing of tangor and sweet orange, but only 661 non-redundant SNP-based DArTseq markers were finally mapped on the integrated map (Curtolo *et al.*, 2017). As shown in Fig. 3 and Fig. 4, several citrus chromosomes have relatively large regions (5-10 Mb) with low recombination, so that many markers in such regions will map as clusters unless the population size is very large. Biological characteristics of citrus, such as long juvenility, seedlessness, ployembryony, apomixis, heterozygosis, gametophytic incompatibility, zygotic selection and gametic selection (Shimada *et al.*, 2014), not only seriously hamper the development of numerous uniquely segregating markers and production of large full-sib populations, but also remarkably influence allelic segregation and recombination ratio (Bernet *et al.*, 2010; Ollitrault *et al.*, 2012a). In addition to parental genotype, differential fitness of gametal genotypes, crossing direction, and regulatory gene interactions likely also contribute to the high level of segregation distortion in citrus (Bernet *et al.*, 2010; Curtolo *et al.*, 2017). In this study, the mapping population are neither full-sib progenies nor interspecific or intraspecific hybrids, but intergeneric hybrids from mixed crosses of several parents. Elimination of polymorphic loci within the parental varieties reduced the number of usable segregating SNP markers. In addition, in comparison to sweet orange, much fewer SNP markers were identified in trifoliate orange. Such problem were also found in all previous trifoliate orange genetic maps (Chen *et al.*, 2008; Lyon, 2008; Ollitrault *et al.*, 2012a). The low availability of segregating markers in trifoliate orange probably due to its low heterozygosis. To maximize detection of segregating SNP markers, greater sequencing depth is needed for the progenies and parents. As the current sequencing depth only resulted in an average coverage of 2.3% of citrus genome, the marker density of our genetic maps could be further improved through increasing the sequencing depth.

Significant differences in linkage group sizes were observed between trifoliate orange and sweet orange genetic maps. Each of the linkage groups of trifoliate orange is larger in genetic length than the corresponding one of sweet orange, even though there were fewer markers mapped on the trifoliate orange genetic map. Similar differences in genetic distances were evident in the previously reported EST-SSR genetic maps of the same parents based on codominant markers segregating in both parents (Chen *et al.*, 2008). Variation in map length was also found between genetic maps of mandarin, pummelo and sweet orange, which all were genotyped with the same set of SNP markers (Ollitrault *et al.*, 2012a). It is impossible to directly compare the genetic distance between specific markers between our genetic maps, because they were separately constructed with markers segregating only in one parent. However, SNP-based genetic linkage maps with high saturation of markers are comparable when referred to physical distance on a reference genome. The variation in genetic map length is unrelated to genome size among different citrus species, which range from 360 to 398 Mb (Gmitter *et al.*, 2012; Wu *et al.*, 2014; Wu *et al.*, 2018). In spite of extensive and significant segregation distortion observed for most of the markers on both maps, such segregation distortion in citrus was proposed to result from gametic selection rather than zygotic selection (Ollitrault *et al.*, 2012a). However, in a simulation study on factors affecting linkage map construction, segregation distortion from gametic selection had little influence on marker order and genetic distance (Hackett and Broadfoot, 2003). Thus, variation of genetic size between linkage group maps of trifoliate orange and sweet orange probably is not due to segregation distortion, but reflects differential recombination rates between the species. Studies on model plants amply demonstrate the impact of genome sequence divergence on recombination rate, and lower recombination rate is related to higher levels of genome divergence (Chetelat *et al.*, 2000; Li *et al.*, 2006a; Opperman *et al.*, 2004). For different citrus-related genera and species, degree of genome heterozygosity differ dramatically. Sweet orange, known to be highly heterozygous, was demonstrated to be a complex interspecific hybrid derived from pummelo and mandarin (Wu *et al.*, 2014; Xu *et al.*, 2013). While trifoliate orange, a genus related to *Citrus* genus, was found to be low in heterozygosity (Chen and Gmitter, 2013). Therefore, in comparison to trifoliate orange, higher heterozygosity in the sweet orange genome probably suppresses recombination frequency, resulting in a smaller genetic size. This is in agreement with the difference in genetic distance of shared markers between genetic maps of Clementine mandarin and pummelo (Ollitrault *et al.*, 2012a). Clementine mandarin is an interspecific hybrid of *C. reticulata* x *C. sinensis* with high genome heterozygosity (Wu *et al.*, 2014), while pummelo is a progenitor species in *Citrus* with low genome heterozygosity (Wang *et al.*, 2017b). In addition, recombination rates are known to differ between sexes in both plants and animals (Lorch, 2005). The size of the male genetic map of Clementine mandarin was notably larger than its female genetic map (Ollitrault *et al.*, 2012a). Our mapping population was a mix of several crosses between different varieties of trifoliate orange and sweet orange with taxa serving as male and female parents in different crosses. However, most of the progenies were generated with trifoliate orange as male parent, which may also contribute to the larger size of the trifoliate orange genetic map.

### Phenotyping and QTL mapping

This is the first report on identification of QTLs related to *C*Las infection and HLB tolerance. Most of the recently reported QTLs in citrus are related to morphological and physiological traits (Asins *et al.*, 2015; Curtolo *et al.*, 2017; Imai *et al.*, 2017; Kepiro and Roose, 2010; Raga *et al.*, 2014; Raga *et al.*, 2016; Sahin-Cevik and Moore, 2012; Sugiyama *et al.*, 2011; Yu *et al.*, 2017; Yu *et al.*, 2016). Only a few reports focused on disease-related QTLs in citrus, such as Citrus tristeza virus (CTV) (Asins *et al.*, 2012; Ohta *et al.*, 2015), Alternaria brown spot (ABS) (Cuenca *et al.*, 2016; Cuenca *et al.*, 2013), and Citrus leprosis virus (CiLV) (Bastianel *et al.*, 2009). Interestingly, only one major QTL was found with considerable effect on the phenotypic resistance for each of these diseases, suggesting that the inheritance of resistance for these diseases is mainly controlled by a single dominant allele. In our study, nine clusters of significant QTLs associated with *C*Las infection were detected on the two parental genetic maps by interval mapping (IM) and restricted multiple QTL mapping (rMQM). In addition, the results of non-parametric Kruskal-Wallis (KW) test were largely in agreement with the IM/rMQM QTL mapping results. Overall, our results suggest that citrus resistance to *C*Las infection is polygenic. In genetic studies, estimated effects of each QTL for disease resistance in plants usually ranges from a few percent to 50 % or more, and a QTL accounting for more than 20% of the phenotypic variation is commonly described as a major or dominant QTL (Davey *et al.*, 2006; St Clair, 2010). Based on these criteria, some of our significant QTLs, respectively located on LG-t7, LG-t8, LG-s3, LG-s5 and LG-s9, would be considered as major QTLs. Thus, we believe that multiple major QTLs are involved in the genetic control of citrus tolerance to *C*Las infection, in contrast to other citrus diseases where resistance can be attributed to a single dominant QTL. Unlike CTV, CiLV and ABS for which strong resistance is available among citrus types, the HLB-suppression of *Poncirus* may be best described as tolerance (Ramadugu *et al.*, 2016). The HLB pathogenetic mechanism is still unknown (Martinelli and Dandekar, 2017; Wang *et al.*, 2017a), and is likely quite complicated. Based on recent progress in studies on HLB, it was proposed that at least three main molecular mechanisms occur in citrus in response to *C*Las infection, and these mechanisms involve many pathways and genes (Martinelli and Dandekar, 2017).

It is noteworthy that these major QTLs were not detected at the same time for all ten time points or the three years, but only a few QTLs were detected at no more than three time points, even though we took the non-significant minor QTLs into consideration. Poor repeatability of QTL detection is a major obstacle in associating QTLs with *C*Las infection in citrus. Generally, Ct value of *C*Las detection was inconsistent among replicate trees and fluctuated widely during the evaluation period, exhibiting large phenotypic variation among samples for most individuals. This was also reported in another long-term field evaluation of *C*Las infection, which focused on a collection of many citrus species and relatives (Ramadugu *et al.*, 2016). Because the spatial distribution of *C*Las within a tree is dramatically uneven, especially at earlier stages of *C*Las infection (Louzada *et al.*, 2016), Ct value, a mean with a high variance, reflects only an average level of *C*Las infection. In a field with HLB pressure, citrus trees are naturally infected by *C*Las through the psyllid vector, and psyllids tend to repeatedly colonize a tree through invading new flush or young leaves (Hall *et al.*, 2016). Moreover, movement of *C*Las occurs slowly from an inoculated position to other branches and finally systemically throughout the whole tree (Albrecht *et al.*, 2014). Other reports where trees were inoculated only at a single time point documented great variability in *C*Las titer over time, with negligible differences between sweet orange and Carrizo (an HLB-tolerant or somewhat-resistant citrange) titers when *C*Las is initially inoculated into leaves, and a tendency for overall decline from initial titer within Carrizo (Stover *et al.*, 2016a; Stover *et al.*, 2016b). These observations suggest that *C*Las titer would be quite dynamic in a field planting where psyllid inoculation is ongoing and sporadic, especially in trees where high titers do not develop. That is why, in spite of continuous natural inoculation with *C*Las for five years, healthy trees could be still occasionally detected within some individuals that had been initially determined to be positive for *C*Las infection.

The phenotypic data quality could be improved by increasing numbers of replicate trees and sampling more frequently or more intensely, however, space requirements and test costs would increase too. Under equivalent conditions, phenotyping with more replicate trees but less sampling repeats is likely more efficient to obtain accurate results in comparison to less replicate trees with more sampling repeats. It is important to note that at least eight replicate trees were clonally propagated for each progeny individual in this study. Although each tree was phenotypically evaluated from a single sample of four leaves at each time point, it is a comparatively cost-effective strategy to ensure better results of QTL mapping. This strategy focused on increasing reliability of mean value for each of the hybrid progenies and parents rather providing accurate estimates of phenotypic traits for each of the trees. Through statistical analysis, we found that the broad sense heritability (*H*^*^2^*^) estimates for Ct value of *C*Las detection range from 0.03 to 0.14 during the evaluation period with 0.09 in average (Supplementary Table 3). The heritability estimates are low, but may be accurate given the many environmental factors that contribute to variability. For example, natural inoculation with *C*Las is related to abundance of psyllid and preference of colonization, as well as the presence and developmental stage of citrus flush (Hall *et al.*, 2016; Lopes *et al.*, 2017). To reduce the influence of non-genetic factors, a more reliable and precise method of *C*Las inoculation is needed. We recently used graft-inoculation with *C*Las on an alternate phenotyping population. Due to limited greenhouse space and the long time commitment for evaluation, we only inoculated 210 trees from 82 of the progeny. Phenotyping results on this graft-inoculated population should improve the robustness of QTL detection.

In conclusion, basing on Genotyping-by-Sequencing of an intergeneric F1 population of 170 progenies, we constructed two high-density genome-wide genetic maps for trifoliate orange and sweet orange. Each of the genetic maps contained nine firm linkage groups corresponding to the haploid chromosome number of the species and exhibited high synteny and high coverage of the reported citrus genome. The minor discrepancies among the genetic maps and genome assembly may represent possible structural rearrangements among citrus species. In replicated field evaluation over three years, trifoliate orange and sweet orange showed significant differences in the Ct value of *C*Las detection and percentage of healthy trees associated with *C*Las infection and tolerance to HLB, and their progenies segregated obviously in these two phenotypic traits. A total of nine clusters of QTLs were found to be associated with *C*Las infection at all time points and years, of which five QTL clusters, respectively located on LG-t7 and LG-t8 of the trifoliate orange genetic map and LG-s3, LG-s5 and LG-s9 of the sweet orange genetic map, collectively explained a major part of the phenotypic variation in response to *C*Las infection. Our results suggest that multiple QTLs are involved in the genetic control of HLB response in citrus. This work provides a starting point for future studies of the underlying genetic architecture of resistance or tolerance to HLB. These QTLs needed to be further confirmed to use this information in breeding for resistance or tolerance to HLB in citrus. In addition, the corresponding genomic regions need to be refined if the objective is to discover and characterize candidate genes related to the disease. The final identified QTLs and genes could be good targets for citrus breeding to support long-term control of this devastating disease.

## Supplementary data

**Table S1** Percentage of healthy trees in control varieties and F_1_ progenies.

**Table S2** Correlation coefficients of phenotypic data.

**Table S3** Broad sense heritability (*H*^*^2^*^) for Ct value of *C*Las detection.

**Fig. S1** Frequency distributions of percentage of healthy trees in F_1_ progenies.

## Acknowledgements

The authors thank Randy Driggers for propagating and planting the trees and Sean Reif for maintenance of the field trees; thank Peng Lin, Bo Yan, Yuan Yu, Misty Holt, Fabieli Lrizarry and Hui Wen for their assistance in sampling. This work was partly supported by grants from the New Varieties Development and Management Corporation (NVDMC), and the Citrus Research and Development Foundation Inc. (CRDF), on behalf of the Florida citrus industry.

